# Epidermal turnover in the planarian *Schmidtea mediterranea* involves basal cell extrusion and intestinal digestion

**DOI:** 10.1101/2023.12.07.567498

**Authors:** Jun-Ru Lee, Tobias Boothe, Clemens Mauksch, Jochen C. Rink

## Abstract

Planarian flatworms undergo continuous internal turnover, wherein old cells are replaced by the division progeny of adult pluripotent stem cells known as neoblasts. How dynamic cell turnover is executed at the organismal scale remains an intriguing question in planarians and biological systems in general. While previous studies have predominantly focused on neoblast proliferation, little is known about the processes that mediate cell loss during tissue homeostasis. Here, we use the planarian epidermis as a model to study the mechanisms of cell removal in *Schmidtea mediterranea*. We established a covalent dye-labeling assay and image analysis pipeline to quantify the cell turnover rate in the planarian epidermis. Our findings indicate that the ventral epidermis is highly dynamic, with a half-life of the constituent cells of approximately 4.5 days. Using live-imaging and pulse-chase assays, we find that epidermal cells undergo internalization via basal extrusion, followed by a migration towards the intestine and ultimately digestion by intestinal phagocytes. Overall, our studies reveal an intricate homeostatic cell clearance process that may reduce the metabolic costs of high turnover tissues in planarians.

## Introduction

Many tissues and organs exist in a state of dynamic turnover, whereby old cells are continuously replaced by the progeny of resident stem cells. For example, in the vertebrate intestinal epithelium, stem cells residing in the crypt continuously generate transit-amplifying cells that differentiate into epithelial cells, which gradually become displaced along the vilus and delaminate once they reach the tip (Barker, 2014; Beumer & Clevers, 2021; Crosnier et al., 2006). The homeostasis of the intestinal epithelium is highly dynamic, with an average life time of epithelial cells on the order of ∼5 days (Tetteh et al., 2016). Besides the precise balancing of cell divisions with cell loss exemplified by the intestinal epithelium, a further conceptual challenge in dynamic tissues is often the process of clearing the cells to be replaced. In the case of liver homeostasis, the resident sinusoidal cells phagocytose apoptotic hepatocytes (Dini et al., 1995; Dini et al., 2002). Similarly, during zebrafish embryonic development, the embryonic ectoderm is capable of phagocytosing dead cells (Hoijman et al., 2021). In other contexts, motile and often non-tissue resident phagocytes mediate cell clearance processes (Ninov et al., 2007). The severe pathological consequences that can result from cell clearance defects underscore the collective importance of the underlying mechanisms (Munoz et al., 2010; Nagata et al., 2010; Savill et al., 2002) and the importance of cell clearance mechanisms in dynamic tissues.

Planarian flatworms provide a fascinating example of dynamic turnover at an organismal scale. Pluripotent adult stem cells, so called neoblasts, are the only division-competent planarian cell type outside of the reproductive system (Cebria et al., 2018; Reddien, 2018; Rink, 2018; Shibata et al., 2010). They are highly abundant at (>10% of all cells) (Baguñà, 2012; Baguna et al., 1989) and reside in the mesenchyme surrounding all internal organs. This implies that all tissue and organ progenitors are generated out of place and thus need to migrate to their target tissues during their terminal differentiation (Atabay et al., 2018; Eisenhoffer et al., 2008; Reddien, 2021). Transplantation of a single pluripotent donor neoblast can rescue the demise of an irradiated host due to the re-colonialization and gradual replacement of host tissues by descendants of the donor stem cell over the course of several months (Wagner et al., 2011), indicating that all tissues are replaced continuously by newly generated neoblast progeny. Finally, planarians do not have a fixed body size. They grow when fed and shrink during starvation, dynamically scaling the total number of organismal cells as a function of feeding history (Baguna & Romero, 1981; Thommen et al., 2019). Overall, the combination of a single source of new cells, organismal cell turnover and body size scaling makes planarians an intriguing model for investigating the underlying principles.

Although the intrinsically dynamic nature of planarian tissue architecture has long been recognized, the field has so far mostly focused on the generation and maturation of neoblast progenitors. The epidermal cell lineage is particularly well studied, due to the disproportionate abundance of epidermal progenitors. Generated by neoblasts expressing the zinc-finger transcription factor zfp-1, newly generated post-mitotic progenitors pass through several differentiation stages during their outward migration and final maturation upon integration in the epidermis (Eisenhoffer et al., 2008; Tu et al., 2015; van Wolfswinkel et al., 2014; Zhu et al., 2015; Zhu & Pearson, 2018). Moreover, the rates of neoblast proliferation and thus likely also progenitor production vary in response to feeding or wounding (Baguna, 1974; Wenemoser et al., 2012). Conversely, much less is known about the modes and mechanisms that govern cell removal during turnover. Apoptotic cell death is known to be regulated in a tissue specific manner (Pellettieri et al., 2010) and cell death processes have been proposed as contributors to de-growth (Gonzalez-Estevez & Salo, 2010). However, neither the mechanisms that induce cell removal nor the processes that mediate cell clearance during turnover are currently known.

In this study, we used the epidermis as a model tissue to explore cell turnover in planarians. We developed a pulse-chase labeling assay to quantify the turnover rate in the planarian epidermis and found that the half-life of ventral epidermis is only approximately 4.5 days. Additionally, we established a live-imaging protocol to visualize the process of epidermal cell removal. Interestingly, we found that epidermal cells were not shed towards the outside, but underwent internalization instead. Furthermore, via dual-color live-imaging and pharmacological treatments, we provide evidence that internalized epidermal cells enter the intestine and are ultimately digested by intestinal phagocytes. Collectively, our results suggest a “march of death” as an integral aspect of epidermal cell turnover and that the self-catabolism in intestinal phagocytes may constitute a general principle in planarian tissue turnover.

## Results

The planarian epidermis is a highly dynamic tissue. Previous BrdU pulse-chase experiments have demonstrated the incorporation of new cells in the planarian epidermis on the time scale of days (Reddien et al., 2005; van Wolfswinkel et al., 2014; Zhu & Pearson, 2018). The rate of turnover, e.g., the half-life of epidermal cells, has not been accurately measured so far. To directly quantify epidermal turnover, we took advantage of the selective accessibility of the epidermis to exogenously supplied substances for the development of a pulse-chase assay. As cartooned in Fig. 1A, our assay relies on the covalent pulse labelling of epidermal cells via a cell-permeant form of the dye Carboxyfluorescein succinimidyl ester (abbreviated as CFSE in the following), followed by the application of a second spectrally distinct covalent label, 7-hydroxy-9H-(1,3-dichloro-9,9-dimethylacridin-2-one) succinimidyl ester (DDAO in the following) after a specific chase interval. Accordingly, changes in the fraction of CFSE^+^/DDAO^+^ versus CFSE^-^/DDAO^+^ cells during the chase period should reflect the fraction of epidermal cells that has undergone replacement during the chase interval. As shown in Fig. 1B, CFSE pulse labelling and immediate fixation results in bright and uniform labelling of the epidermis under our experimental conditions. The dye labelling pattern in cross sections (Fig. 1C) indicates labelling specificity for the most superficial cell layer, i.e., the epidermis. Similar results were observed for DDAO labelling (not shown). Hence CFSE and DDAO selectively label epidermal cells without any signal interference from the deeper tissue layers. As expected, the fraction of CFSE^+^/DDAO^+^ cells indeed decreased with increasing time separation between the CFSE and DDAO pulses (Fig. 1D), which is consistent with the gradual replacement of the cells present during the initial pulse with newly incorporated, CFSE^-^ cells (Fig. 1D).

**Figure 1.**
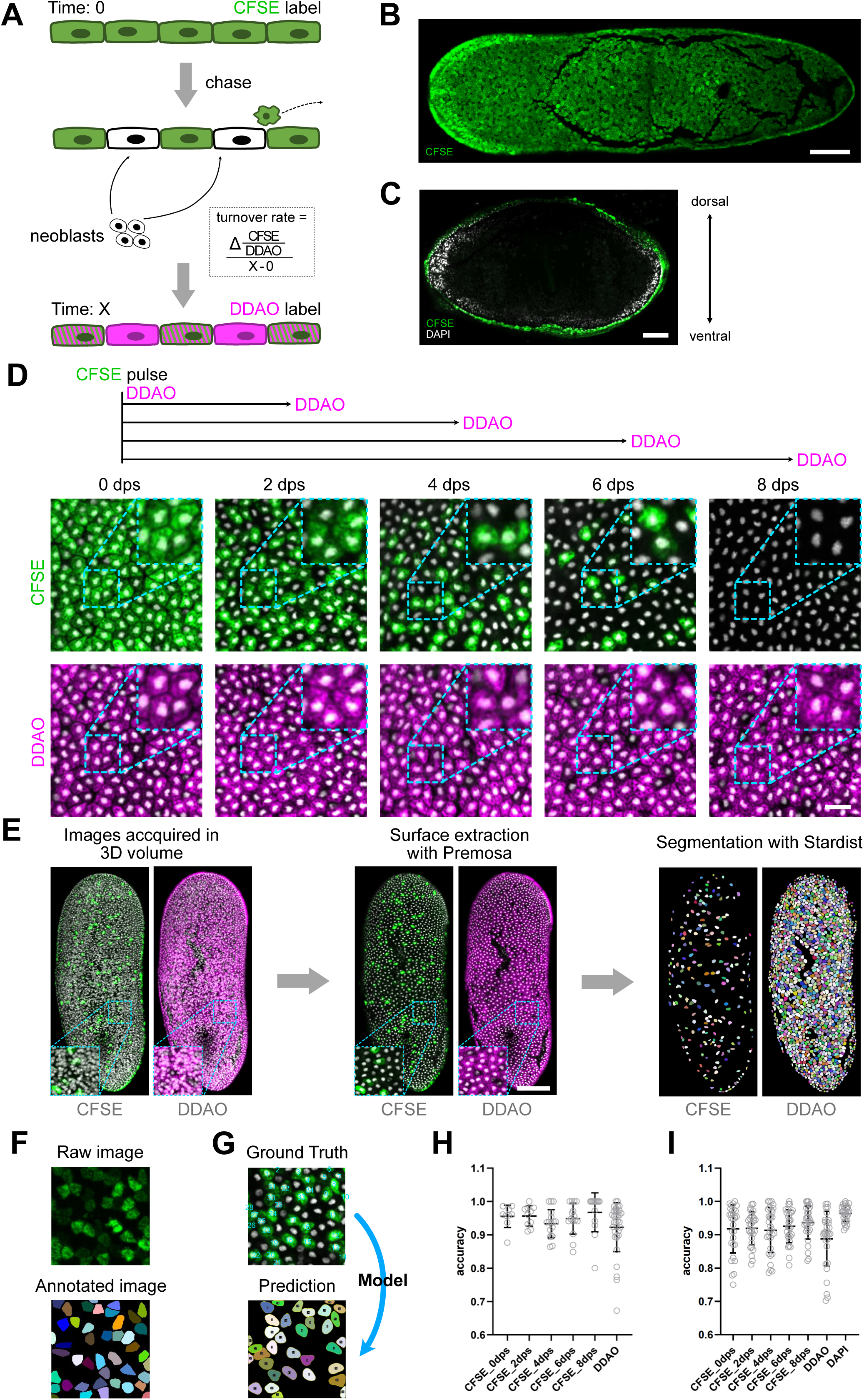
Establishment of an assay for the quantification of epidermal cell turnover. **A**: Schematic of the labeling assay for the quantification of epidermal cell turnover, see text for details. Green: Initial CFSE pulse label; Magenta: DDAO pulse label at assayed time point. The fraction of CFSE^+^ amongst all DDAO^+^ cells/assay time interval represents the runover rate. **B**: Surface-extracted image of the CFSE pulse-labelled ventral epidermis. Cracks in the epidermis (black lines) are a mounting artefact, but illustrate the epidermal specificity of the label. Scale bar: 100 μm. **C:** Sagittal cross-section after CFSE pulse-labelling, further demonstrating the epidermal specificity of CFSE labelling. Scale bar: 100 μm. **D:** Assay time series schematic (top), showing the gradual loss of CFSE+ cells (green, top image panels) versus DDAO^+^ cells (bottom image panels) within the ventral epidermis. DAPI-counter staining of epidermal nuclei is shown in gray. Zooms highlight the decreasing fraction of CFSE^+^ cells. dps: days post CFSE staining. Scale bar: 20 μm. **E:** Schematic of the image analysis procedure for the quantification of CFSE^+^ and DDAO^+^ cells, comprising 3D volume acquisition (left panel, maximum projection), 2D epidermal surface extraction (center panel) and automatic segmentation of epidermal cells (right panel). Scale bar: 100 μm. **F-G:** Manual annotations (F) were used as ground truth training data for segmentation model training and refinement (G). **H-I:** Accuracy assessment of the trained Stardist segmentation model in the ventral (H) and dorsal (I) epidermis at the indicated time points post CFSE labeling. 1 corresponds to equal cell counts in automatically segmented images versus the corresponding manually annotated ground truth. Each open circle represents an individual cell count comparison. N = 100 analyzed cell fields in H; n = 210 in I. Error bar: standard error. Central line: mean.

Towards the goal of quantifying the labelled fractions and thus the turnover rate in our pulse-chase assays, we combined volume-imaging via spinning disk confocal microscopy (necessary due to the intrinsic curvature of the planarian epidermis) with the establishment of an image analysis pipeline (Fig. 1E). The surface extraction tool Premosa was used to generate a 2D representation of the epidermis from the 3D volumes (Blasse et al., 2017), in which individual epidermal cells were subsequently segmented via the deep learning-based object segmentation method Stardist (Schmidt et al., 2018) (See materials and methods for Stardist model training and performance estimation). Using manually annotated datasets of altogether 27668 cells as ground truth, the Stardist model achieved a segmentation accuracy exceeding 90% for both CFSE and DDAO positive cells of the ventral epidermis (Fig. 1H). In the dorsal epidermis, the accuracy of the Stardist model in CFSE labelled epidermal cells also exceeded 90% (Fig. 1I), but the detection of cells in the DDAO channel was less reliable (accuracy ≦ 90%). We therefore used the quantification of DAPI-positive nuclei in the extracted epidermal layer as alternative total cell count (accuracy > 90%) (Fig. 1I). Overall, our image analysis pipeline allows for the precise quantification of old and new cells in the planarian epidermis and thus the quantification of turnover rate of the tissue.

With the assay at hand, we first set out to determine the turnover kinetics of the multi-ciliated ventral epidermis, which planarians rely on for their gliding motility (Vu et al., 2019). Quantifications of the ratio of CFSE^+^/DDAO^+^ cells over time indicated that on average, 80% of the epidermal cells underwent replacement within 8 days (Fig. 2A). On basis of these data, we calculated the half-life of epidermal cells (i.e., the chase time by which 50% of the originally labelled cells have been replaced) as 4.5 d (Fig. 2C). Importantly, animal cohorts assayed 7d post irradiation displayed much slower and gradually flattening kinetics of CFSE^+^ cell loss (Supp. Fig. 1). This result is consistent with the depletion kinetics of epidermal progenitors post irradiation (Tu et al., 2015) and thus confirms the assumption that our assay measures cell replacement rather than heterogeneities in label retention. Given the distinct morphological features and patterning signal dependencies of the ventral and dorsal epidermis in planarians (Kato et al., 2001; Molina et al., 2007; Reddien et al., 2007; Rompolas et al., 2010; Wurtzel et al., 2017) we also assayed the turnover rate of the dorsal epidermis. Interestingly, we found that the turnover rate in the dorsal epidermis was much slower than in the ventral epidermis, with ∼80% of the original cell cohort remaining 8 days after the initial pulse (Fig. 2B). Direct comparison of the turnover rates between dorsal and ventral epidermis via linear regression analysis revealed a half-life of ∼20 days dorsally as compared to the ∼4.5 days ventrally (Fig. 2C). Our results directly demonstrate tissue specificity in turnover rates and that the ventral epidermis is an especially dynamic tissue.

**Figure 2.**
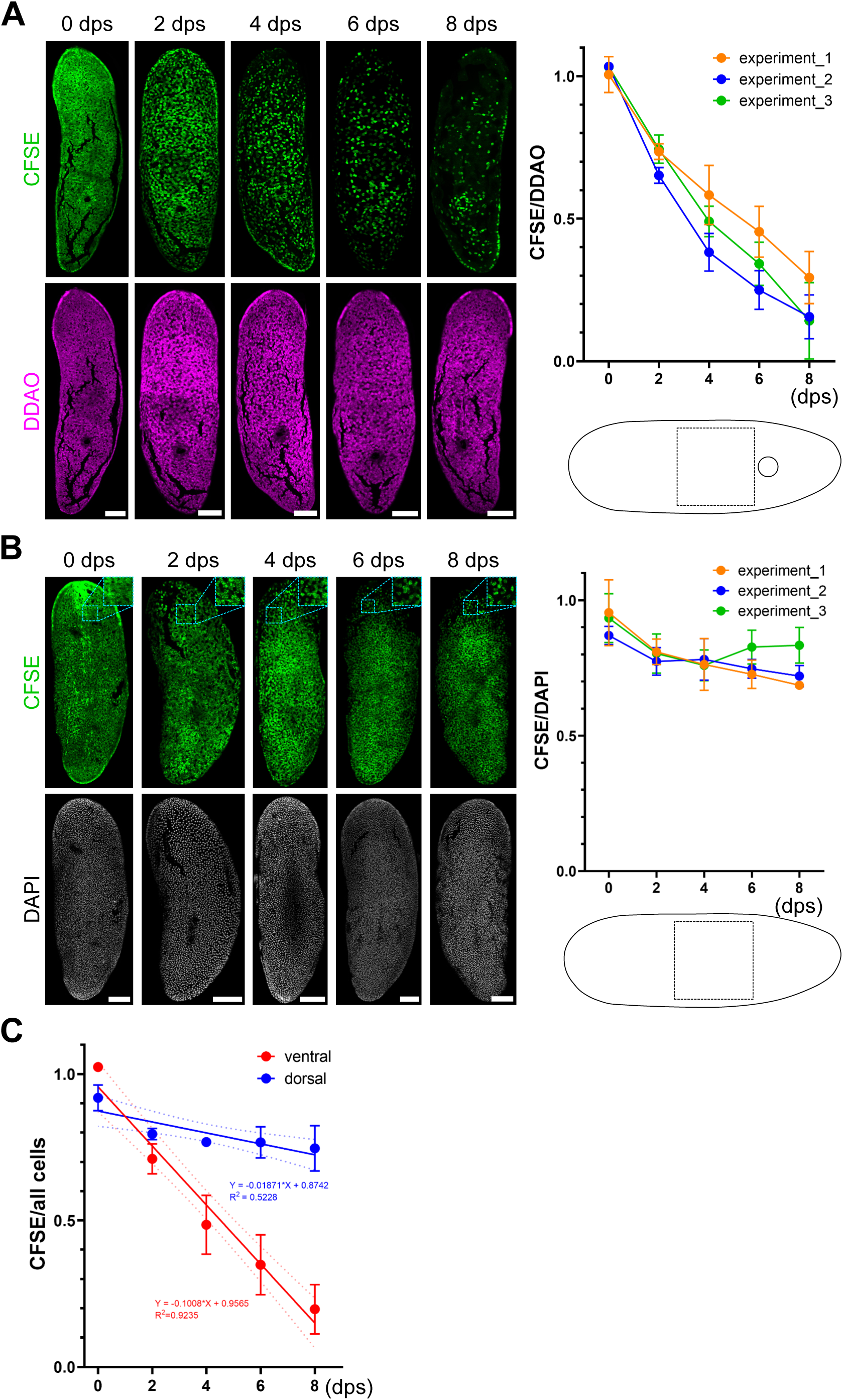
Turnover kinetics quantification of the planarian epidermis. **A-B**: Quantification of the cell turnover rates in the ventral (A) and dorsal (B) epidermis. Left: Representative specimens stained with the indicated probe combination at the indicated days post staining. Right: Quantifications of the CFSE^+^ cell fraction within she square-shaped analysis area anterior to the ventral mouth opening (circle) as diagrammed. N = 3 biological replicates (experiments), each comprising 3-6 specimens/time point. Error bars denote the standard error. Scale bars: 100 μm. **C:** Linear regression analysis of the turnover rates in ventral and dorsal epidermis, based on the data in A-B. The dashed lines represent the 95% confidence intervals, error bars represent the standard error between replicate means.

In principle, epidermal cells can either undergo shedding towards the outside (apical extrusion, as in the vertebrate intestine) or undergo basal extrusion (delamination towards the inside). To distinguish between these possibilities, we thought to live-image epidermal turnover. Live imaging planarians remains a challenge and planarian cell dynamics have so far not been visualized. Building on our previous efforts (Boothe et al., 2017; Boothe et al., 2023), we developed a live imaging procedure involving the combination of Linalool anaesthetization and agarose embedding under a gas permeable Polymethylpentene (thermoplastic polyolefin) disc to immobilize the worms. Besides, low-excitation imaging of far-red live dyes was implemented to minimize the strong photosensitivity of planarians especially to shorter wavelengths (Shettigar et al., 2017) and post-acquisition AI-denoising (Fig. 3A). Specifically, we used the nuclear dye RedDot™1 and the cytoplasmic label CellTracker™ Deep Red, which were both brighter and more photostable than DDAO under our imaging conditions (not shown). Despite these measures, occasional contractions of the body wall musculature (twitches in the following) still interfered with the intended cell tracking. As shown in Fig. 3B, a further reduction of excitation intensity and exposure times to 5 ms (see Material and Methods) and post-acquisition drift correction to correct the remaining twitches enabled the stable tracing of RedDot™1-stained epidermal nuclei for up to 2 hours (Fig. 3B). CARE-mediated de-noising (Weigert et al., 2018) of the low signal to noise image sequence further achieved sufficiently accurate nuclear detection for long-term tracking and for addressing the direction of cell extrusion in the planarian epidermis.

**Figure 3.**
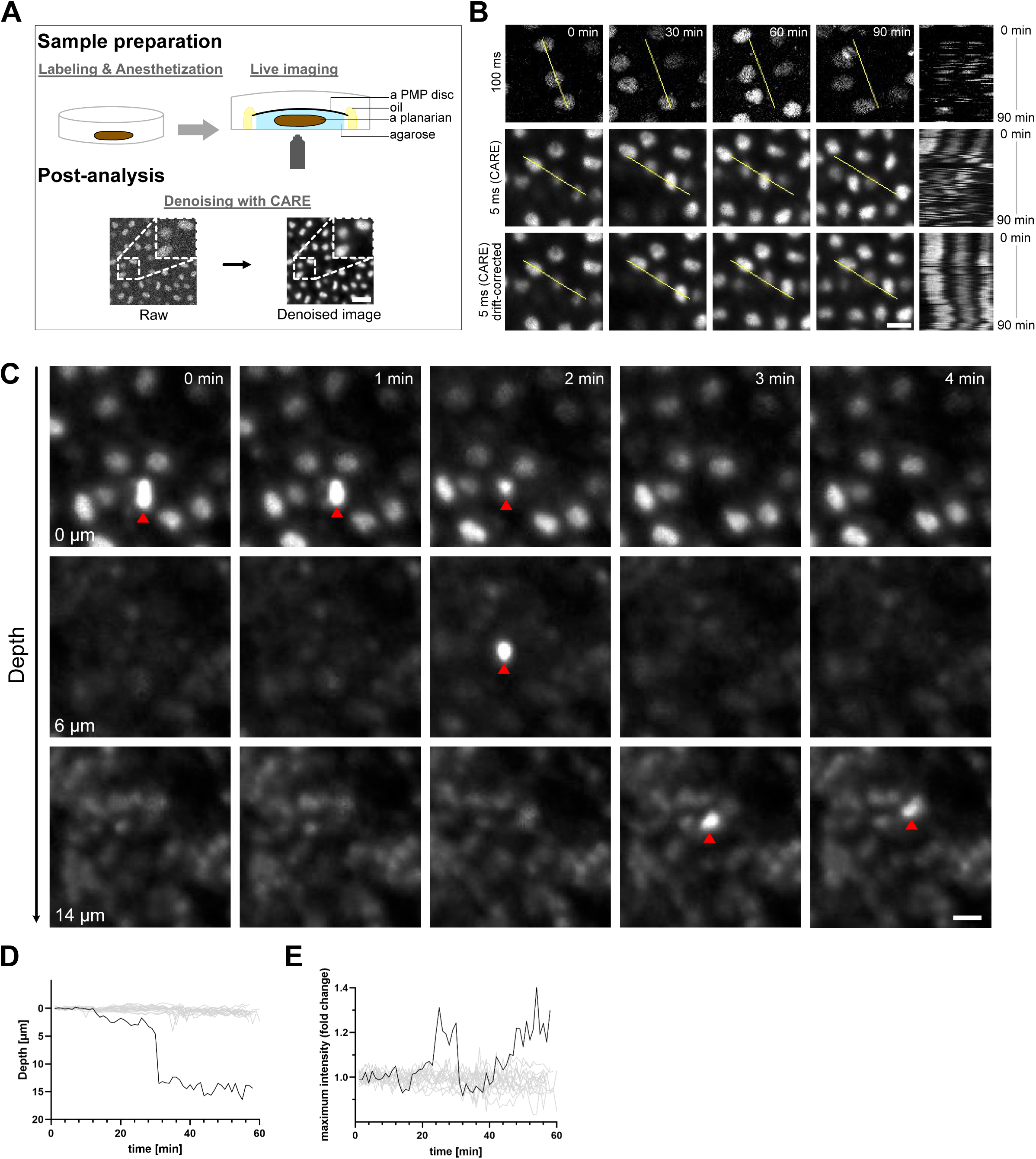
Live-imaging of epidermal cell turnover. **A:** Cartoon of the live-imaging procedure, involving Linalool anaesthetization, mounting in low-melt agarose underneath a PMP disc, spinning disc confocal Z-stack time lapse imaging and post-processing. See text for details. Scale bar: 20 μm. **B**: Live imaging optimizations of epidermal cells labelled with the far-red fluorescent nuclear probe RedDot™1. Rows show a representative crop of a timelapse image sequence at the indicated time points. The right column displays kymographs of pixel brightness over time along the yellow line in the respective image panel. Top row: 100 milliseconds exposure time/Z-stack image. Center row: 5 milliseconds exposure time/Z-stack image and CARE de-nosing. Bottom row: 5 milliseconds exposure time/Z-stack image, CARE de-nosing and additional drift correction. Scale bar: 10 μm. **C**: Internalization of a RedDot™1-labelled epidermal nucleus. The image frames trace the same cell field in individual confocal sections (5 msec exposure time, CARE de-nosing, drift correction) at the indicated time points (left to right) and the indicated Z-depths (top-bottom). The red triangle highlights the internalizing nucleus. Scale bar: 10 μm. **D-E**: Quantification of the relative z-depth (D) or maximum pixel intensity (E) of an internalizing nucleus over time (black line), relative to non-internalizing epidermal nuclei (gray lines).

As expected from a terminally differentiated barrier epidermis, the epidermal nuclei in our time-lapse sequences were mostly static and were thus easily detectable under our imaging conditions. Interestingly, in 6 post-processed movie sequences spanning an approximate total of 12 hours and 4800 cells, we never observed the extrusion of an epidermal nucleus towards the outside. Instead, we observed nuclei undergoing rapid vertical migration towards deeper tissue layers, traversing a zone mostly devoid of nuclei (Fig. 3C, Supp. Movie 1) prior to encountering more densely populated layers. The nuclear movements typically occurred over a vertical distance of ≧ 15 um (Fig. 3C, D). Live-imaging with a cytoplasmic label (CellTracker™ Deep Red) provided further indicated that the vertical movements represent basal extrusion events with gap closure by the surrounding cells (Supp. Movie 2). Epidermal cell internalization events were comparatively rare at 1.5 events per 2 h with approximately 800 nuclei per field of view, which is somewhat lower than the event frequency expectation on basis of the measured t1/2 of 4.5 days (see discussion). Overall, our live imaging experiments therefore indicate that the turnover of the planarian epidermis involves the basal extrusion and further internalization of epidermal cells.

In light of these results, the further fate and eventual destination of the internalized epidermal cells became an important question. As a first approach, we applied two-color live imaging of epidermal nuclei together with prior feeding of a fluorescent dye to label the intestine as a landmark of deep tissues. Interestingly, we could observe that the basally extruded epidermal cells rapidly descend into the close proximity of intestinal branches (Fig. 4A). The walls of the planarian intestine largely consist of so-called phagocytes, which are known to actively phagocytose material via their surface (Ishii & Sakurai, 1991; Willier et al., 1925). The maximally achievable live imaging duration of 2 h was insufficient for ascertaining the further fate of internalized nuclei. As an alternative approach, we returned to our CFSE pulse-chase assay and asked whether the CFSE surface label might eventually reach the intestine. Worms fixed immediately after CFSE staining were entirely devoid of CFSE fluorescence in all sub-epidermal tissues, thus again confirming the epidermal specificity of the labelling strategy (Fig. 4B, 4D). However, 4-days post CFSE staining, CFSE^+^ objects were clearly visible in deep tissues, where they partially colocalized with phagocytosed CellTracker™ Deep Red delivered by feeding 1 day prior to fixation (Fig. 4B). Moreover, quantifications of the CFSE^+^ object density within the volume occupied by the CellTracker™ Deep Red label versus the mesenchyme (see materials and methods for details) indicated that the epidermis-derived structures specifically accumulate within gut branches (Fig 4C, 4D). To unequivocally demonstrate the accumulation of CFSE^+^ structures inside phagocytes, we combined CFSE pulse-chase with antibody signal amplification (see Methods) and *in situ* hybridization with a phagocyte-specific probe (*dd_00075*; (Fincher et al., 2018)). As shown in Fig. 4E, CFSE^+^ structures were indeed detected within the cytoplasm of phagocytes after 4 days of chase, but not immediately after labeling. Overall, our results suggest that epidermal cells are continuously internalized to accumulate in intestinal phagocytes.

**Figure 4.**
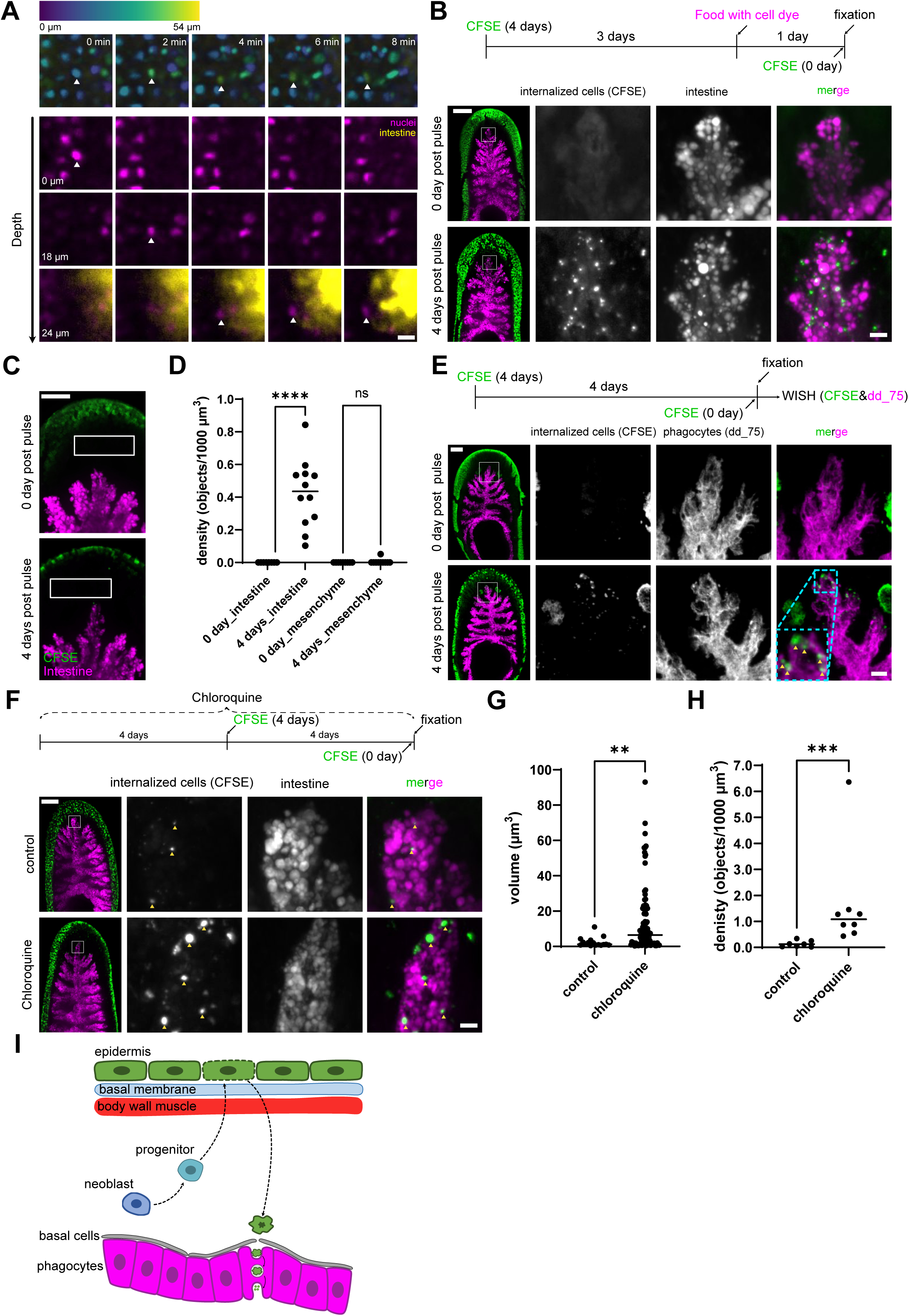
Tracing the fate of basally extruded epidermal cells. **A**: Two-color time-lapse imaging of RedDot™1-labelled epidermal cells and CellTracker™ Red CMTPX Dye-labelled intestinal cells. Top row: Depth-coded maximum-projected images of the of RedDot™1 channel at the indicated time points. The rows below display two channel merged images at the indicated z-depth. Channel colour coding as indicated. White arrowheads: internalizing epidermal cell. Imaging conditions: Simultaneous/sequential acquisition of the two channels, 5 msec exposure times and post-processing CARE de-noising and drift-correction. Scale bar: 10 μm. **B**: The CFSE-surface label can be chased into the intestine. Top: Cartoon representation of the experiment. Detail images reveals CFSE-fluorescence within intestinal branches at 4 days post staining (bottom), but not 0 dps (top). White frames in the overview (left) designate the location of the detail image. All images are maximum-projections. Channel color coding as indicated. Scale bars overview images: 100 μm; Scale bar detail images: 10 μm. **C, D**: Quantification of CFSE^+^ object density within and outside the intestine under the same assay conditions as in C. CFSE^+^ objects and the intestinal area were identified using the “Threshold” function in FIJI. The “mesenchyme” analysis box (white outlines in C) was positioned between the epidermis and intestine. The number of CFSE^+^ objects and volume of intestines and mesenchyme were quantified using Cellprofiler after Fiji thresholding. 8-12 worms were analyzed in each experimental condition. Each solid circle corresponds to the quantification from one individual worm. The central line represents the median. The Mann-Whitney U-test was used to compare the density differences. ****: p<0.0001; ns: non-significant. Scale bar: 50 μm. **E**: Confirmation of intestinal CFSE accumulation by antibody staining (anti-Fluorescein) and in situ hybridization with the phagocyte-specific probe *dd_75_0-1*. Top: Cartoon of the experiment. Detail images and zooms displays anti-Fluorescein staining within phagocytes at 4 days (bottom), but not 0 days of chase (top). Yellow arrowheads: CFSE^+^ objects. White frames in the overview (left) designate the location of the detail image. All images are Maximum-projections. Channel color coding as indicated. Scale bar (left panel): 100 μm; Scale bar (right panel): 20 μm. **F**: Accumulation of internalized CFSE label in the presence of the lysosome inhibitor Chloroquine. Top: Cartoon of the experiment. Detail images highlight the increased size and number of intestinal CFSE^+^ structures (yellow arrowheads) in the presence of Chloroquine (bottom) versus control (top). The white frame in the overview image to the left designates the location of the detail image. All images are Maximum-projections. Channel color coding as indicated. Scale bar (left panel): 100 μm; Scale bar (right panel): 10 μm. **G-H**: Quantifications of intestinal CFSE^+^ object volumes (G) and density (H) under the same experimental conditions as in F. CFSE^+^ objects and intestines were identified using the “Threshold” function in FIJI. The volume and overall density of CFSE^+^ objects and the intestine were quantified using Cellprofiler after Fiji thresholding. N = 7 worms for control, N = 8 for the chloroquine treatments. Solid circle indicates individual CFSE^+^ objects in G or the overall density within one sample in H. Central lines represent the median. The Mann-Whitney U-test was used to assess statistical significance. **: p<0.01 ***: p<0.001. **I:** Model of epidermal turn-over in planarians, involving a “march of death” of internalized epidermal cells. See discussion for details.

This raises the possibility that phagocytes might not only catabolize ingested food, but also endogenous cells undergoing turnover. Accordingly, CFSE^+^ remnants of internalized epidermal cells should accumulate in the intestine under experimental conditions that interfere with its digestive functions. We therefore treated planarians with the lysosome inhibitor Chloroquine, which blocks intracellular digestion by interfering with the acidification of late endosomal compartments (Halcrow et al., 2021; Homewood et al., 1972; Mauthe et al., 2018; Schlesinger et al., 1988). Chloroquine exposure had no overt effects on planarians with the tested concentration, even when applied for more than 1 week. Interestingly, we found that Chloroquine exposure indeed strongly increased both the volume and also the density of intestinal CFSE^+^ objects above controls cultured in plain planarian water (Fig. 4F-H). Both observations are consistent with reduced digestion of phagosomal contents in the presence of Chloroquine. Moreover, Chloroquine treatment also increased the density of CFSE^+^ objects in the mesenchyme 4 d post CFSE treatment (but not at t0; Fig. S3), indicating that Chloroquine might additionally affect the uptake of internalized epidermal cells into phagocytes. Similarly, interference with phagocyte differentiation and function via knock-down of *hnf4*, a transcription factor that is expressed in the intestinal lineage (van Wolfswinkel et al., 2014; Wagner et al., 2011), resulted in the accumulation of CFSE^+^ structures in the mesenchyme (Fig. S2), consistent with the clearing of epidermal remnants by phagocytes. Overall, our results therefore identify the digestion of internalized epithelial cells by intestinal phagocytes as the last stage of the complex turnover of the planarian epidermis.

## Discussion

Using new pulse-chase and live imaging assays, we here address the kinetics and physiological mechanisms of cell turnover in the planarian epidermis. We find that the ventral epidermis is a highly dynamic tissue with a half-life of only 4.5 days and that surprisingly, cell turnover is mediated via internalization, rather than shedding. As cartooned in Fig. 4I, our results collectively reveal that epidermal cells undergo a “march of death”, involving the crossing of the thick basement membrane underneath the epidermis, the 3 layers of the body wall musculature and a section of mesenchyme to become phagocytosed and eventually digested by the phagocytes that make up the walls of the intestine. Besides representing an unusually complex sequence of events for the clearing of aged cells, the “march of death” also retraces in reverse the outward migration that every epidermal progenitor undergoes after its “birth” in an asymmetric cell division of a pluripotent neoblast situated deep within the mesenchyme (Eisenhoffer et al., 2008; Tu et al., 2015; van Wolfswinkel et al., 2014). With long-range cell migration as part of both cell birth and death, our results add an additional element to the already extraordinarily dynamic tissue architecture of planarians.

Our live imaging results cannot rule out that the “march of death” is not the only turnover mechanism of epidermal cells. Assuming desynchronised and stochastic turnover kinetics at steady state, the measured t1/2 of ventral epidermal cells of 4.5 days corresponds to an hourly internalisation probability of 4.5 cells per 800 cells. The observed internalisation frequency of ∼ 0.75 events/∼ 800 cells/h under our imaging conditions is therefore ∼ 6 times lower than expected. Similarly, we never observed new cell integration in any of our time lapse sequences, which should occur at approximately the same frequency as internalisation events. Possible explanations for this discrepancy could include anaesthetization or dye labelling side effects or adverse effects of our mounting/imaging conditions, which will need to be addressed in future studies. Nevertheless, the combination of internalisation as the only observed turnover mode of epidermal cells (Fig. 3C), the live tracing of individual nuclei down to the vicinity of intestinal branches (Fig. 4A), the delayed appearance of the covalent CFSE surface label in intestinal phagocytes (Fig. 4B) and the confirmation of signal specificity by antibody enhancement (Fig. 4E) or the effects of Chloroquine treatment (Fig. 4F-H) collectively support the digestion of internalized epidermal cells in intestinal phagocytes as last stage of their “march of death”.

As depicted, the “march of death” envisages active migration of internalized epidermal cells down to the intestine. Due to their flat body architecture and elaborate branching of the intestine, we estimate that epidermal cells even in large planarians have to maximally traverse a distance of <100 um even in large planarians to reach the closest intestinal branch, which is entirely within range of cell migrations in other systems (Ninov et al., 2007; van Ham et al., 2012). Moreover, the fact that we could track individual nuclei from the epidermis all the way down into close proximity of the intestine and the short time duration of the migration (∼6 min) support the hypothesis that the march of death reflects active and autonomous cell migration. However, our present data cannot rule out less parsimonious transfer mechanisms, for example the participation of migratory phagocytic cells in the transfer of internalized epidermal cells to intestine.

The catabolism of aged epidermal cells in a distant organ is unusually elaborate in comparison with clearing mechanisms in other model systems. Most animal epithelia shed cells towards the outside, with the constitutive shedding of vertebrate intestinal epithelium cells at the cristae tip (Heath, 1996; Potten, 1992, 1997) or the apical extrusion of epidermal cells in the tail of developing zebrafish (Eisenhoffer et al., 2012) as examples. Although basal extrusion is also known to exist in other systems (Ninov et al., 2007; Pastor-Pareja et al., 2004; Toyama et al., 2008), it is more frequently associated with pathologies with basal extrusion and consequent metastasis formation upon oncogenic k-Ras transformation as one example (Marshall et al., 2011; Slattum et al., 2014; Slattum & Rosenblatt, 2014). The crossing of the collagen-rich basement membrane (Chan et al., 2021; Cote et al., 2019; Hori, 1979) that planarian epidermal cells undertake post basal extrusion establishes an interesting conceptual parallel to metastasis formation in vertebrates, but also raises the question regarding the eventual purpose of the march of death in planarians.

Although entirely speculative at the time being, the self-catabolism of epidermal cells may serve as an energy recovery mechanism. Planarians never reach a stable body size. They grow when fed, but shrink during periods of starvation, gradually down-scaling all organs as a function of the steady decrease of the number of the total cell count (Baguna & Romero, 1981; Thommen et al., 2019). The starvation response is termed “de-growth” and it has long been speculated that self-catabolism satisfies the energy demands of planarians during starvation. Our results likely provide a first glimpse of the underlying processes. The quantitative considerations above illustrate that even the self-catabolism of one of the most short-lived cells types results in a comparatively low retrograde cell flow at steady state. It is therefore entirely plausible that epidermal cells are not the only planarian cell type that end their existence in a “march of death” and that the intricate branching of the planarian intestine facilitates both, the distribution of externally acquired nutrients, as well the efficient “recycling” of constituting cells. Which additional cell types undergo a march of death, the turnover rates of different organs and whether or not they are regulated during growth and de-growth constitute just some of the interesting questions for future exploration. In addition, our results raise a number of new cell biological questions. One intriguing problem are the triggering mechanisms that initiate basal extrusion and ultimately determine the half-life of epidermal cells and its tissue specificity (Fig. 2, 3C). The concomitant brightness increase and condensation of epidermal nuclei prior to internalization (Fig. 3E) may hint at a participation of apoptotic processes, as in basal extrusion in the *Drosophila* amnioserosa or larval epidermis (Levayer et al., 2016; Nakajima et al., 2011; Ohsawa et al., 2018; Teng et al., 2017; Toyama et al., 2008). The downward migration of internalized cells likely requires migration tracks (e.g., D/V muscle fibers) or guidance cues for directional migration that remain to be identified. A further important question is whether phagocytes phagocytose epidermal cell remnants on the mesenchymal side or from the luminal side subsequent to paracellular transfer into the intestinal lumen. Of note, live-imaging approaches will be necessary in understanding those aspects of planarian cell biology. Our study provides important proof of principle in this respect and the approaches that we have pioneered are a starting point for the systematic investigation of this dynamic dimension of planarian biology.

## Acknowledgements

We thank Maria Lina Moroni for generously sharing the DAPI-labeled cross sections, and Ludwik Gasiorowski for his critical reading of the manuscript draft. This work was supported by the Max Planck Society and the Behrens-Weise-Foundation.

## Author contributions

J.-R.L., T.B. & J.C.R conceptualized the project and designed the experiments. J.-R.L., T.B. & C.M. conducted the experiments. J.-R.L. & T.B. analyzed the results. T.B. provided the technical supports for the image analysis methods. J.-R.L., T.B. & J.C.R wrote the manuscript.

## Materials and Methods

Key reagents and tools table

| Reagent or Source | Source | Identifier |
| --- | --- | --- |
| **Animal model** |  |  |
| *Schmidtea mediterranea* | Grown in our lab | NA |
| **Antibodies** |  |  |
| Anti-DIG-POD | Roche | 11207733910 |
| Anti-FITC-POD | Merck | 11426346910 |
| **Chemicals & Reagent** |  |  |
| CellTracker™ Deep Red Dye | Invitrogen | C34565 |
| CellTracker™ Red CMTPX Dye | Invitrogen | C34552 |
| CellTrace™ CFSE Cell Proliferation Kit | Life Technologies | 6585084 |
| CellTrace Far Red DDAO-SE | Invitrogen | C34553 |
| Chloroquine Diphosphate Salt | Sigma-Aldrich | C6628 |
| Ciprofloxacin | Sigma-Aldrich | 17850 |
| DAPI | Thermo Scientific | 62247 |
| DIG RNA labeling mix | Roche | 11277073910 |
| Gibson Assembly Master Mix | New England BioLabs | E2621S |
| Halocarbon oil 700 | Sigma-Aldrich | H8898 |
| Linalool | Sigma-Aldrich | L2602 |
| N-Acetyl-L-cysteine | Sigma-Aldrich | A7250 |
| Paraformaldehyde 16% solution, EM grade | Electron Microscopy Sciences | 15710 |
| Reddot1™ Far-Red Nuclaer Stain, 200X in water | Biotium | 40060 |
| SeaPlaque™ Agarose | Lonza | 50100 |
| **Software** |  |  |
| Affinity Designer | Serif Ltd | NA |
| CARE | (Weigert et al., 2018) | NA |
| CellProfiler | (Carpenter et al., 2006) | NA |
| FIJI | (Schindelin et al., 2012) | NA |
| IMARIS | Oxford Instruments | NA |
| Premosa | (Blasse et al., 2017) | NA |
| Prism | GraphPad Software | NA |
| Stardist | (Schmidt et al., 2018) | NA |
| **Others** |  |  |
| CellRad Precision Irradiator | Precision X-RAY irradiation | NA |
| μ-Dish 35mm | ibidi | 81158 |

### Animal culture

All experiments were performed on the asexual strain of *Schmidtea Mediterranea* (CIW4). The animals were cultured in planarian water (1.6 mM NaCl, 1.0 mM CaCl_2_, 1.0 mM MgSO_4_, 0.1 mM MgCl_2_, 0.1 mM KCl and 1.2 mM NaHCO_3_ in Milli-Q water) containing 5 μg/mL Ciprofloxacin in plastic boxes at 20 °C. The worms were fed with macerated calf livers once every two weeks. Animals were last fed 1 week or 2 weeks before the experiments depending on the designs of the experiments. Ciprofloxacin was removed from waste water prior to disposal via charcoal filtration.

### Irradiation

2 mm worms were selected and irradiated with X-ray at a dosage of 60,000 rad using the CellRad Precision Irradiator. The worms were rinsed twice with Planarian water containing 5 μg/mL Ciprofloxacin immediately after irradiation exposure and the planarian water was then refreshed daily.

### Cloning and Probe synthesis

The target genes were amplified from *Schmidtea mediterranea* cDNA with gene-specific primer pairs and inserted into the pPRT4P vectors using Gibson assembly. The plasmid constructs were used for *in situ* hybridization probe synthesis or double-stranded RNA synthesis later. Linear DNA amplified by PCR from the gene-inserted plasmids were used as the templates for anti-sense probe synthesis. *In vitro* transcription for probe synthesis was performed by incubating 1 μg DNA template, 1X in vitro transcription buffer (25 mM MgCl_2_, 40 mM Tris pH 8.0, 10 mM DTT, 2 mM spermidine), 1X DIG RNA labeling mix (Roche, Cat 11277073910), 1 U/μL RNAse inhibitor (Thermo Scientific, Cat EO0384), 0.0015 U/mL Inorganic Pyrophosphatase (Thermo Scientific, Cat EF0221) and 2 U/μL T7 polymerase (Thermo Scientific, Cat EP0111) together at 37 °C overnight. The probe RNA was then precipitated by mixing the *in vitro* transcription mixture (20 μL) with 10 μL 7.5 mM ammonium acetate and 50 μL ice-cold 100% EtOH. After centrifugation, the RNA pellets were resuspended in deionized Formamide (AppliChem, Cat A2156)

**Table 1.**
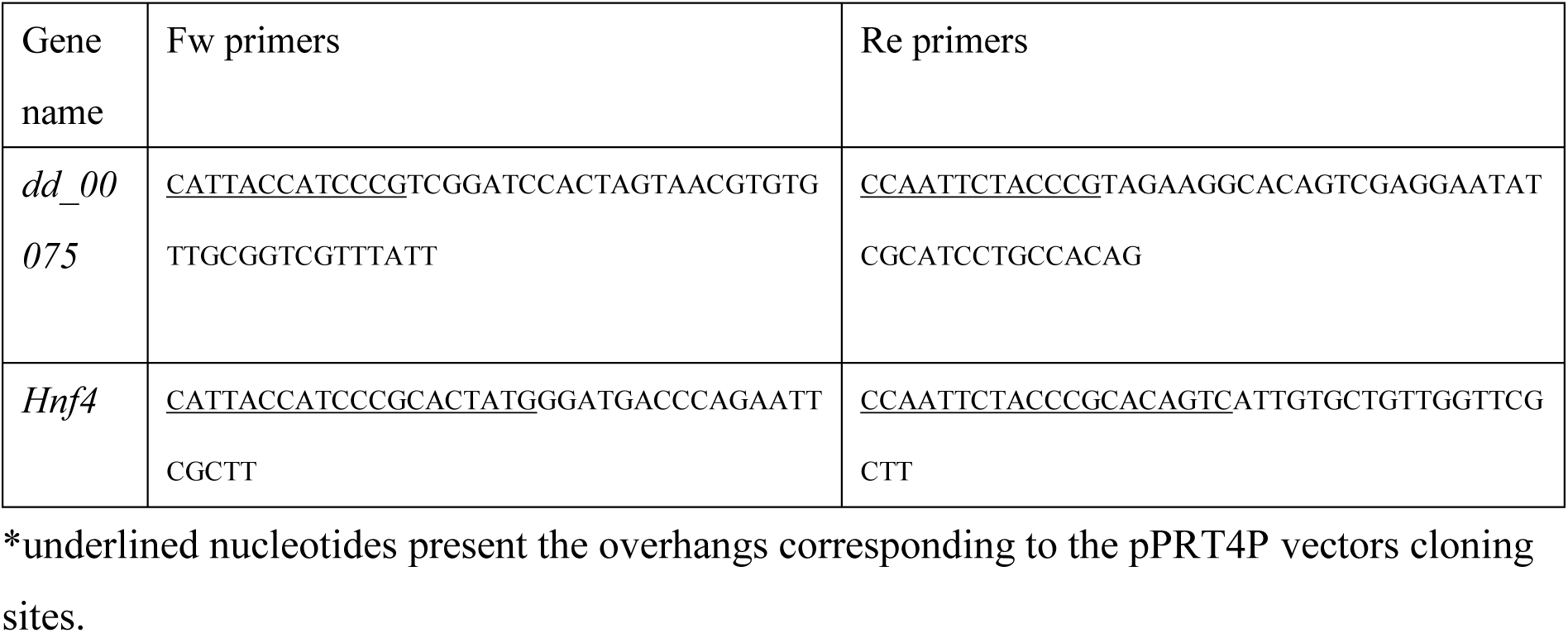
Primer pairs used for probe generation.

### RNA interference

In vitro transcription for double-stranded RNA synthesis was conducted by incubating 1 μg DNA template, 1X in vitro transcription buffer (25 mM MgCl_2_, 0.04 M Tris pH 8.0, 0.01 M DTT, 2 mM spermidine), 7.5 mM rNTP, 1 U/μL RNAse inhibitor (ThermoFisher, Cat EO0384) and 3.4 U/μL T7 polymerase (ThermoFisher, Cat EP0111) together at 37 °C overnight. After *in vitro* transcription, single-strand RNA was annealed by heating at 75 °C and gradually cooling down to room temperature. The annealed RNA was then precipitated using NaCl/PEG-8000 precipitation solution (2.5 M NaCl, 20% Polyethyene glycol 8000, 10 mM Tris-HCl, pH 8.0) through centrifuging at 4 °C. The target gene was then silenced by feeding the double-stranded RNA (dsRNA) mixed with calf liver paste to the animals. The final concentration of the dsRNA was 2 mg/mL. Worms were fed with dsRNA twice a week. For *hnf4* RNAi, worms were fed three times in total before the further experimental operations.

### Whole-mount *in situ* hybridization

Whole-mount *in situ* hybridization was conducted as described previously in (King & Newmark, 2013) with a few modifications: In brief, worms were euthanized in 5% NAC in PBS, fixed in 4% PFA in 50% PBS containing 0.15% Triton-X. Worms were soaked in reduction solution (1% NP-40, 0.5% SDS, 50 mM DTT in 1XPBS in H_2_O) at 37°C for 10 minutes and dehydrated in 100% MeOH. After rehydration in PBS containing 0.3% Triton-X, worms were bleached in bleaching solution (1.2% H_2_O_2_, 5% formamide, 0.5XSSC in H_2_O) on a light table for 1 hour and treated with 2 μg/mL Protease K (NEB, Cat PB107S) in PBS containing 0.3% Triton-X for 10 minutes followed by post-fixation in 4% PFA for 10 minutes. The samples were soaked in Pre Hybe (50% Formamide, 5X SSC, 1X Denhardts, 100 μg/μL Heparin, 1% Tween-20, 1 mg/mL torula yeast RNA, 50 mM DTT) at 58 °C for 2 hours and were incubated in Hybe (50% Formamide, 5X SSC, 1X Denhardts, 100 μg/μL Heparin, 1% Tween-20, 0.25 mg/mL torula yeast RNA, 50 mM DTT, 0.05 g/mL dextran sulfate) with RNA probes at 58 °C overnight. The next day, samples were washed with Wash Hybe (50% Formamide, 0.5% Tween-20, 5XSSC, 1 X Denhardts) twice; 1:1 Wash Hybe: 2XSSC (0.1% tween-20) twice; 2XSSC (0.1% tween-20) three times; 0.2XSSC (0.1% tween-20) three times at 58 °C. Samples were soaked in blocking solution (5% sterile horse serum, 0.5% Roche western blocking reagent in TNTx) for two hours at room temperature. Following blocking, samples were incubated with antibody (anti-DIG-POD or anti-FITC-POD) in blocking solution (1:1000 from the antibody stocks) at 4 °C overnight. For fluorophore development, samples were washed with TNTx (0.1 M Tris pH 7.5, 0.15 M NaCl, 0.3% Triton X-100 in H_2_O) six times and proceed to tyramide amplification by incubating in TSA buffter (2M NaCl, 0.1M Boric acid in H_2_O, pH8.5) containing 0.006% H_2_O_2_, 20 μg/mL 4-Iodophenylboronic acid and tyramide (Rhodamine or FAM) for 45 minutes at room temperature. For CFSE-labeled worms, CFSE signal was amplified by incubating in anti-FITC-POD antibodies (1:1000) in blocking solution and the fluorophore was developed by tyramide amplification (Rhodamine). The peroxidase activity of first antibody was quenched by incubating in 100 mM sodium azide in TNTx for at least one hour at room temperature. The color reaction of the second antibody was developed as the first one. After color development, samples were soaked in Scale S4 (10% Glycerol, 15% DMSO, 40% Sorbitol, 4 M Urea, 2.5% DABCO, 0.1% TritonX-100 in H_2_O) at 4 °C overnight and mounted on slides the next day.

### Paraffin sections with DAPI labeling

The sectioning procedure followed the published protocol (Solana, 2018). Initially, 2 mm worms were euthanized in cold 2% HCl in planarian water and fixed in 4% PFA at room temperature for 30 min under slow rotation. This was followed by an overnight wash in PBS at 4°C. The samples were then dehydrated in ethanol, followed by washes in xylene and were eventually embedded in clean paraffin. The embedded worms were incubated in paraffin for 1 hour on a cold plate until the paraffin solidified followed by at 4 °C overnight incubation. The solidified samples were then cut into 3 μm thick sections and incubated at 37 °C until they looked smooth. These sections were mounted on poly-L-lysine coated glass slides and incubated at 37 °C overnight. Nuclei were labeled with 0.5 μg/mL DAPI in PBS for 30 min at room temperature. Finally, the slides were mounted in ProLong ^TM^ Glass Antifade Mountant (Thermo, Cat P35984).

### CFSE and DDAO labeling for epidermis

The epidermis was labelled with CFSE and DDAO lively by immersing the worms in planarian water containing 10 μM of CFSE or 1 μM of DDAO for 1.5 hours at 20°C. The worms were then euthanized in 5 % N-Acetyl-L-cysteine in PBS for 3 minutes on a rotator and fixed in 100% MeOH at −20°C for 2 hours. Following fixation, the samples were rehydrated sequentially with a series of MeOH concentrations in PBS: 75% MeOH, 50% MeOH and 25% MeOH. Nuclei were stained by immersing the worms in 5 µg/mL DAPI or 5 µg/mL Hoechst in PBS for 30 minutes at room temperature followed by three PBS rinses. Lastly, the samples were stored in 80% glycerol in PBS for at least 30 minutes at room temperate before being mounted between two of coverslips (24*60 mm, No. 1.5H and 22*22 mm, 1.5H; Schott) to allow for imaging from dorsal and ventral sides. For the cross-section samples, the specimens were mounted in 4% SeaPlaque^TM^ agarose and cut into two pieces using a razor knife.

### Live-imaging for the planarians

The nuclei of epidermis were lively labeled with 2x Reddot™ 1 and 1% DMSO in planarian water at 20 °C overnight. For the cytoplasm labelling, worms were immersed in planarian water containing 10 µM CellTracker™ Deep Red Dye at 20 °C for 1.5 hours. To label the intestine of the planarian, worms were fed with liver paste containing fluorescent dye mixture (20% liver paste, 0.3% SeaPlaque^TM^ Agarose, 0.02 mM CellTracker™ Red CMTPX Dye in water) adopted form (Forsthoefel et al., 2011) one day prior to the time-series imaging. Following labeling, the worms were anesthetized in 0.02% linalool at room temperature for 30 minutes. Mucus was removed by immersing worms in neutralized (approx. pH 7) 0.5% N-Acetyl-L-cysteine containing 0.02% linalool in distilled water for 10 seconds. Finally, worms were transferred to a μ-Dish 35 mm with a glass bottom and embedded in 1% SeaPlaque^TM^ agarose containing 0.02% linalool. The agarose and sample were then covered with a PMP plastic disc. The gap between the cover and glass bottom was sealed with halocarbon oil 700.

### Image collection

The fixed samples were imaged using an Olympus IX83 equipped with a Yokogawa CSUW1-T2S Spinning Disk system, which was fitted with a Hamamatsu Orca Flash4.0 V3 camera. Imaging was conducted with a 20X air objective (NA = 0.8). The samples were scanned with 1 μm section spacing.

For live animal samples, imaging was collected on an Olympus IX83 microscope paired with a Yokogawa CSUW1-T2S Spinning Disk system and an Andor iXon Ultra 888 EMCCD camera. A 60X Silicone objective (NA = 1.3) or a 30X Silicone objective (NA = 1.05) was used. The samples were incubated in a thermostatic chamber at 15°C for the whole process of time-series imaging. Prior to capturing the time series images, animals were incubated in the chamber for 30 minutes. 640 nm and 561 nm excitation lasers were used for imaging with a laser power of 1 % and an exposure time of 5 milliseconds. The samples were scanned over a total distance of 30-60 μm with 1 or 2 μm each section. Z-stacks were recorded at one-minute intervals over a period of three hours.

### Image processing

#### Premosa surface extraction

The surface cells (epidermis) were extracted from the 3D images of CFSE, DDAO and DAPI-labeled samples with Premosa software (Weigert et al., 2018). DDAO signals which labeled all epidermal cells served as the reference for extracting CFSE signals and nucleus signals from the surfaces.

#### Stardist model training

To train Stardist models, raw images and manually annotated images served as the input training pairs for model trainings. The raw images were the Premosa-processed CFSE-labeled images. For the ventral epidermis, both images from un-irradiated and irradiated worms were used. 0.3% pixels of the images acquired from the un-irradiated worms were normalized to saturation and 256*256 pixel crops were randomly selected. For the images acquired from irradiated worms, crops were made followed by 0.3% pixels normalized to saturation. For training images in dorsal epidermis, 1% pixels of the images from the unirradiated worms were normalized to reach saturation and the 256*256 pixel crops were randomly selected. These crops served as the raw images for training. The annotated images came from the manual annotation of individual epidermal cells from the raw images with Labkit in FIJI. Additionally, each raw image and annotated image were rotated by 90, 180, 270 degrees and every image including the original and rotated ones was flipped. Therefore, 8 different raw and annotated images were generated from one original image by the rotation and flip. For the training of Stardist models, a total of 199*8 training pairs were used for training the ventral epidermis model, and 66*8 pairs were used for training the dorsal epidermis model. The detailed parameters for the training of the Stardist models are listed below.

**Table 2.** Training parameters for Stardist models.

|  |  |  |
| --- | --- | --- |
| Training Parameters | Number of rays (n_rays) | 64 |
|  | Number of patch size (train_patch_size) | 128, 128 |
|  | Number of epochs (train_epochs) | 200 |
|  | Steps per epoch | 200 |
|  | Batch size (train_batch_size) | 32 |
|  | Learning_rate (train_learning_rate) | 0.0003 |

#### Stardist model verification

For the Stardist model verification, 256*256 pixel crops were selected randomly in DDAO, CFSE and DAPI labeled images. Prior to cropping, 1% or 3% of pixels were normalized to saturation in CFSE or DDAO labeled images. The detailed parameters for the Stardist model prediction are listed below. In the ventral epidermis, a 200 pixel^2^ was set as the size cutoff to eliminate the oversegmented objects. The results from the model prediction were compared to the results from the ground truth (manual counting). The accuracy of the Stardist models was quantified using the following equation:

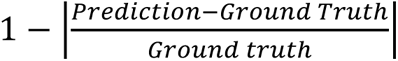. For the turnover rate quantification, the same parameters as employed in the model verification were applied to the selected area.

**Table 3.** Parameters for the quantification of CFSE^+^, DDAO^+^ and DAPI^+^ objects with Stardist.

|  | Ventral epidermis |  | Dorsal epidermis |  |  |
| --- | --- | --- | --- | --- | --- |
|  | CFSE | DDAO | CFSE | DDAO | DAPI |
| Models | Custom-trained ventral epidermis model | Custom-trained ventral epidermis model | Custom-trained dorsal epidermis model | Custom-trained dorsal epidermis model | Built-in fluorescent nuclei |
| % pixels to saturation | 1 | 3 | 1 | 3 | 0 |
| Neural Network Prediction |  |  |  |  |  |
| Percentile Low | 0 | 0 | 0 | 0 | 0 |
| Percentile High | 100 | 100 | 100 | 100 | 100 |

| NMS Postprocessing |  |  |  |  |  |
| --- | --- | --- | --- | --- | --- |
| Scored threshold | 0.6 | 0.6 | 0.4 | 0.4 | 0.6 |
| Overlap threshold | 0.3 | 0.3 | 0.3 | 0.3 | 0.3 |

#### CARE model training

To enhance the signal-to-noise ratio of time-series images, CARE models were trained using z-stack images captured consecutively with low exposure time (5 milliseconds) and high exposure time (100 milliseconds). Two to five image pairs were used for a single training. The parameters for training CARE models are listed below.

**Table 4.**
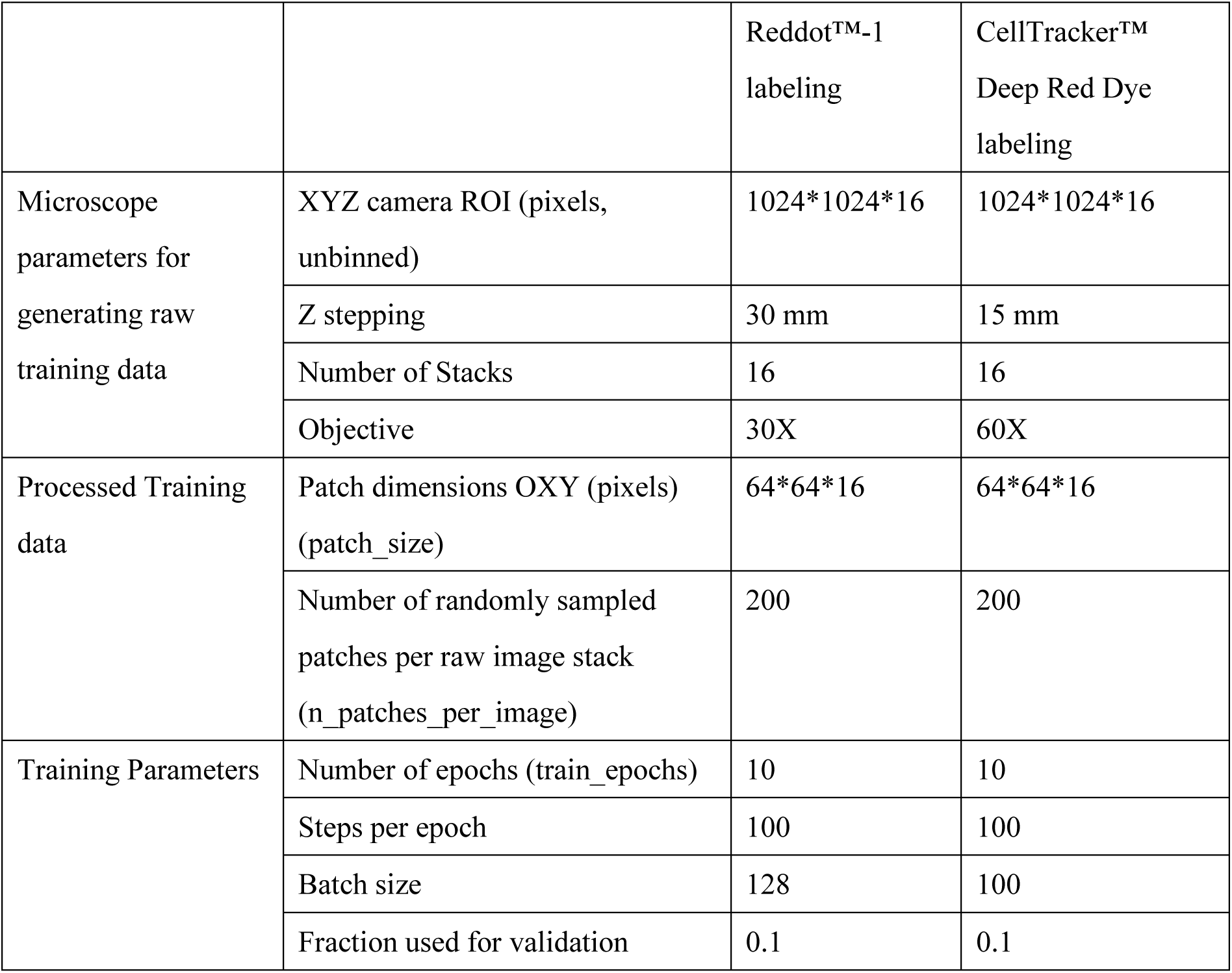
Parameters for the CARE model trainings.

#### Visualization of time-series images

For the visualization of time-series images, the specific regions were selected and the signal-to-noise ratio was enhanced using the previously trained CARE models. To correct the drift of the sample over time, linear stack alignment with SWIFT in FIJI was used to align the images from single-color live-imaging recordings. Fast4DReg was used for the correction of two-color live-imaging. The deepness of the z-stack images was represented with different colors by the built-in color LUT (mpl-viridis) in FIJI.

#### Quantification of CFSE^+^ volume and density

To quantify the intestine volume, each CFSE^+^ object volume and the number CFSE^+^ objects, the gut branches in the anterior dorsal tip ends were chosen. Greyscale images were transformed to binary masks with the threshold function in FIJI. The Huang model was used for thresholding. The binary images were subjected to analysis with the CellProfiler pipelines. The CFSE^+^ objects that are smaller than 10 voxels were eliminated. To quantify the ratio of the CFSE^+^ volume to total volume of the selected area, the mesenchymal between the intestine and epidermis in anterior region were selected. The volume was quantified with the CellProfiler pipelines.

#### Tracking of epidermal cells in time-lapse recordings

Cropped, CARE-restored and drift corrected z-stacks were processed with IMARIS software for tracking individual nuclei over time. In brief, objects were detected 3D by intensity and size thresholding. The IMARIS-implemented object tracking function was used to track nuclei over time and analyzed their relative position and intensity throughout the time course.

#### Drug treatment

For chloroquine treatment, 3 mm worms were immersed in 5 μM Chloroquine in planarian water containing 5 μg/mL Ciprofloxacin for 4 days and the epidermis was labeled with CFSE for 1 hour on the fifth day of chloroquine treatment. The control worms (0 day-post labeling) were fixed in 100 % MeOH immediately after the CFSE pulse. The remaining worms were fixed 4 days after the CFSE labeling. Throughout the CFSE chase, worms were kept in planarian water with 5 μM Chloroquine. To visualize the intestine, worms were fed with liver paste containing fluorescent dye (20% liver paste, 0.3% SeaPlaque^TM^ Agarose, 0.02 mM CellTracker™ Deep Red Dye in water).

#### Statistics

The Mann-Whitney-U test was used for all of the analyses conducted in this study.

