## Supplementary figures and images for "Epidermal turnover in the planarian *Schmidtea mediterranea* involves basal cell extrusion and intestinal digestion"

### Supplemental Figures

Supplemental Figure 1

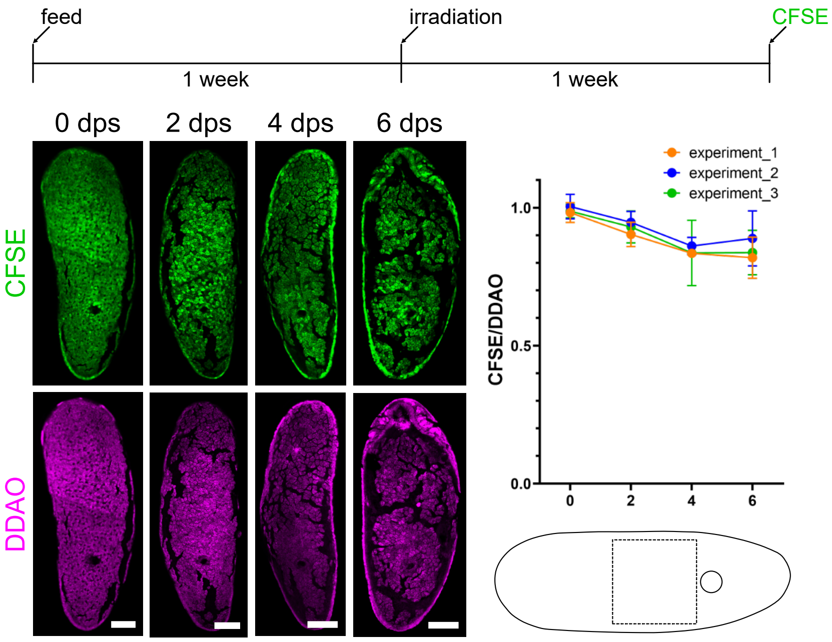

Supplemental Figure 2

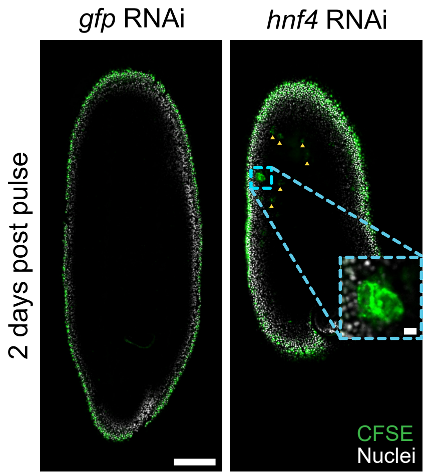

Supplemental Figure 3

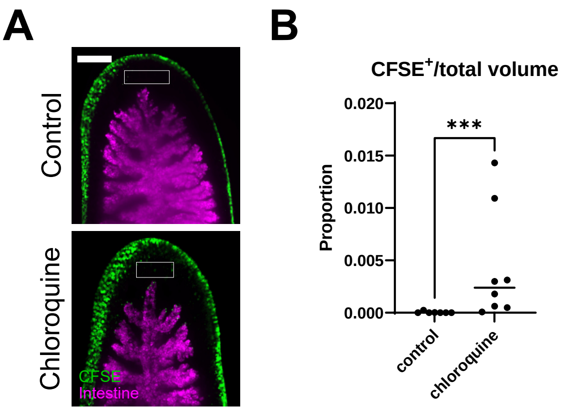
